# Association of NQO1 C609T (Pro187Ser) with risk of Oral Submucous Fibrosis in Eastern Indian population

**DOI:** 10.1101/046052

**Authors:** Sanjit Mukherjee, Sweta Mohanty, Atul Katarkar, Richa Dhariwal, Basudev Mahato, Jay Gopal Ray, Keya Chaudhuri

**Affiliations:** CSIR-Indian Institute of Chemical Biology, Kolkata, India; Department of Oral Pathology, Dr. R. Ahmed Dental College & Hospital, Kolkata, India

**Keywords:** Areca nut, Oral submucous fibrosis, NQO1, single nucleotide polymorphism, carcinogenesis

## Abstract

Oral submucous fibrosis (OSF) is a debilitating disease mainly attributed to chewing areca nut with a 7.4-13% malignant transformation rate. Present study explores the role of NADPH quinone oxidoreductase 1 (NQO1) C609T (*Pro*187*Ser*) polymorphism in susceptibility to OSF among habitual areca nut chewers in an eastern Indian population. Overall, about 18% of the total OSF cases were detected carrying minor TT allele (*Ser/Ser*) p=0.026. When categorized by age, both CT (*Pro/Ser*) and TT (*Ser/Ser*) alleles were significantly higher (p= 0.003 & 0.004 respectively) in cases above 40years of age. NQO1 protein was 42% reduced in buccal tissues of heterozygous (*Pro/Ser*) carriers, whereas a 70% reduction was observed in TT (*Ser/Ser*) OSF cases. Our study suggests that the NQO1 C609T polymorphism confers increased risk for OSF in habitual chewers.

## Introduction

Upsurge in the popularity of commercially prepared areca nut preparations (popularly known as *‘pan-masala & gutka’*), as a chewing habit have contributed to high prevalence of oral carcinogenesis in India (Rajendran, 1994) (Ray et al., 2019) (More et al., 2020). Chewing of areca nut with betel quid increases the concentrations of carcinogenic nitrosamines and reactive oxygen species in mouth (Nair et al., 1992) (Adil et al., 2021). As an early sign of damage to oral mucosa, chewers often develop precancerous oral lesions such as leukoplakia and submucous fibrosis. Oral submucous fibrosis (OSF) is an insidious, chronic, progressive precancerous condition of the oral cavity and oropharynx with a high degree of malignant potentiality. A significant number of this precancerous condition converts into Oral Squamous Cell Carcinoma (OSCC), the rate being about 7.4-13% (Aziz, 1997). This disease is now a public health concern in many parts of the world including many southeast Asian countries, United Kingdom and South Africa though it is mainly prevalent in the Indian subcontinent in all age groups and across all socioeconomic strata (Gupta and Warnakulasuriya, 2002) (Warnakulasuriya et al., 2002). The human NAD(P)H:quinone oxidoreductase 1 gene (*NQO1*; DT-diaphorase, Enzyme Commission (EC) number 1.6.99.2) occupies 17 kilobase pairs (kb) within a gene-rich region on chromosome 16 at 16q22.1 (Jaiswal et al., 1988). This cytosolic flavoenzyme detoxifies quinones (a large class of aromatic compounds found commonly in plants, benzene metabolites, and chemotherapies) to hydroquinones or catechols. The enzyme NAD(P)H:quinone oxidoreductase 1 (NQO1) acts as an antioxidant by catalyzing a 2-electron reduction that bypasses the need for two 1-electron reductions that can result in the production of DNA and protein-damaging reactive oxygen species. In certain conditions (e.g., the presence of myeloperoxidase or autooxidants), NQO1 can contribute to the formation of reactive oxidation species via oxidative cycling and therefore can act as a prooxidant (Ross et al., 2000). NQO1 is constitutively expressed in most tissues including the bone marrow, where expression is thought to be highly inducible by xenobiotics with quinone moieties and is upregulated during times of oxidative or electrophilic stress. The polymorphisms of NQO1gene have been characterized and known for about two decades (Traver et al., 1997). A C>T substitution at position 609 of *NQO1* cDNA, which codes for a Proline > Serine (*Pro>Ser*) change at residue 187 has been found to be associated with risk of several cancers till date(Lajin and Alachkar, 2013). Functionally, a rapid degradation of NQO1 was observed in colon cancer cells transfected with a mutant 609T vector *in vitro*. Presence of a *Pro>Ser* mutation rapidly targets the NQO1 protein for ubiquitin mediated proteasomal degradation pathway leading to reduced NQO1 activity in these cells (Siegel et al., 2001). The prevalence of the minor T allele varies among ethnic groups (4–20%), with the highest prevalence occurring in Asian populations (Kelsey et al., 1997; Gaedigk et al., 1998). In one of our previous observational studies we found a very high prevalence of NQO1 *Pro>Ser* polymorphism among both urban and rural population of West Bengal, India, who were habitually exposed to processed arecanut and smokeless tobacco (Ray et al., 2013). This study explores the role played by NQO1 *Pro>Ser* polymorphism in increasing susceptibility to Oral Submucous fibrosis among habitual areca nut and smokeless tobacco chewers.

## Materials and Methods

### Patients and controls

A total of 179 newly diagnosed cases of oral submucous fibrosis and 152 healthy control subjects having oral habit for a similar period were recruited from outpatient’s department at Dr. R Ahmed Dental College & hospital, Kolkata, India over a 3-year period (2010-2013). All individuals enrolled in the study provided informed consent for their participation in the study. All participants completed a questionnaire providing information about their age, gender, ethnic origin, and use of processed areca nut products, smoking or alcohol consumption status. A written consent was obtained from all the patients after educating them about the importance and outcome of the study. Blood was collected from the antecubital vein, while biopsy sample was collected from the affected tissue. All the study protocols were reviewed by an institutional ethical committee, and samples were collected by medical trained professionals. A part of the tissue was used for routine histopathological and immunohistochemical examination, while another part was used for further analysis of genetic or protein expression.

### Polymorphism Genotyping

Genotyping was carried out by PCR-RFLP analysis on DNA extracted from whole blood samples obtained from all participants in the study using primers forward: 5’AAGCCCAGACCAACTTCT-3’ and reverse:5’-ATTTGAATTCGGGCGTCTGCTG-3’ with an initial denaturation of 95^0^C for 2 mins followed by 35 cycles of 94°C for 30 secs, 60°C for 30secs, 72°C for 1 min and a final elongation of 72°C for 10 mins. Subsequently, the PCR products were digested with 20 U of HinfI (New England Biolabs, USA) for 3 hr at 37°C and separated on a 6% polyacrylamide gel. The NQO1 wildtype allele shows a 172 bp PCR product resistant to enzyme digestion, whereas the null allele shows a 131 bp and a 41 bp band. The frequency of the NQO1 genotypes was compared in the patient and control groups.

### RNA isolation from oral mucosal tissue

Total RNA was extracted using Trizol reagent (Gibco BRL, Gaithersburg, MD) according to the manufacturer’s protocol. Approximately 50 mg of tissue was collected in 1 ml Trizol reagent and homogenized using a handheld homogenizer. After the addition of 0.2 ml chloroform, the mixture was vigorously shaken for 3 min at 22°C and centrifuged at 13000 rpm for 15 min at 4°C. An equal volume of isopropanol (chilled) was added and was kept in cold condition for 10 min, RNA was precipitated by centrifugation at 13000 rpm for 10 min at 4°C. The pellet was washed twice with 70% ethanol, briefly dried under air, and dissolved in 100 µl of diethylprocarbonate-treated water.

### Quantitative realtime PCR of NQO1 mRNA

Isolated RNA from OSF and control tissues was immediately subjected to realtime RT PCR (using Takara PrimescriptTM one step realtime RT PCR kit) (Takara, Japan) analysis for detection of changes in expression of NQO1 mRNA according to the manufacturers protocol. Briefly, the PCR was carried out using following primers Forward 5’-GGGCAAGTCCATCCCAACTG-3’ and Reverse 5′GCAAGTCAGGGAAGCCTGGA-3’ (230 bp) while for GAPDH the forward and reverse primers were 5’-ATGGGGAAGGTGAAGGTCGG-3’ and 5’-GGATGCTAAGCAGTTGGT-3’ respectively, yielding a 470 bp product. The reaction mix contained 200 ng of RNA; PrimeScript One Step Enzyme Mix; 1X One Step Buffer; 20 µM of each of the primers and RNase Free dH2O. The real-time RT-PCR cycling program involved an initial denaturation step at 95°C for 2 min, followed by 40 cycles of 15 s at 95°C and 30 s at 60°C. Thermal cycling was performed in a BioRad iQ5 Real-Time PCR Detection System (Hercules, CA, USA).

### Western blotting

The tissues were collected in PBS buffer containing protease inhibitor cocktail (Sigma-Aldrich, USA). Tissue extracts were prepared by RIPA lysis buffer [150 mM NaCl 1% Nonidet P-40 (vol/vol) 0.5% AB-deoxycholate (vol/vol) 0.1% SDS (vol/vol) 50mMTriszHCl (pH 8) 1mM DTT 1mg/ml each of leupeptin, aprotinin, and pepstatin]. The insoluble pellet was discarded, and the protein concentration was determined by using Bradford reagent (Bio-Rad). Equal amounts of protein were mixed with sample buffer (4% SDS 20% glycerol 10% 2-mercaptoethanol 0.125 M TrisHCl), heated at 95**°**C for 5 min, loaded and separated on 6% polyacrylamide-SDS gel. After separation, the proteins were blotted onto a PVDF membrane (Millipore, Billerica MA) and incubated with mouse monoclonal NQO1 (1:1000) (Cell Signalling Technology, Beverly, MA) and β-Actin Antibody (C4) (1:2000) (Santa Cruz Biotechnology, USA) as primary antibodies. Goat anti mouse secondary antibody conjugated with ALP (1:2000 dilution) was then added and incubated for 3 hours at room temperature followed by washing in TBST six times for ten minutes each to detect the protein levels. The alkaline phosphatase-positive bands were visualized using a 5-bromo,4-chloro,3-indolylphosphate/nitrobluetetrazolium (BCIP/NBT) (GENEI). The band’s intensities were analyzed using ImageJ (http://rsbweb.nih.gov/ij/download.html) and expressed in densitometric units (DU).

### Immunohistochemistry

Paraffin blocks each from wild type, heterozygous and homozygous mutant OSF cases were selected for subsequent immunohistochemical investigations. Briefly, dissected oral tissues from biopsy specimen were paraffin embedded and 4-5µm thickness sections were collected on poly-L-lysine coated slides, after paraffin removal by xylene and rehydration, the slides were treated with citrate buffer for unmasking the antigen. The further immunostaining was performed using Novolink polymer detection system (Leica Microsystems, Switzerland). The endogenous peroxidase and protein were blocked using supplied blockers. The expression of NQO1 protein were detected using NQO1 monoclonal antibody (Cell signaling technologies, Beverly, MA). After post-primary blocking the sections were incubated with Novolink polymer and were then developed with DAB using supplied substrate buffer. The sections were then counterstained with haematoxylin and were observed under LEICA DM 3000 microscope ((Leica Microsystems, Switzerland).

### NQO1 activity assay from affected and adjacent normal tissue

Oral biopsy tissues were obtained from cases (OSF and OSF associated with malignancy) both from the lesion and from the tissue adjacent to the lesion for measurements of NQO1 enzyme activity. The assay was performed as described by Ernster (1967) (Ernster and Ronald W. Estabrook, 1967). and modified by Benson et al. (1980) (Benson et al., 1980) using DCPIP as a substrate. The reaction mix contained 25 mM Tris (pH 7.4), with or without 0.07% bovine serum albumin (w/v) (Sigma-Aldrich, USA) as activator, 200µM NADH (Sigma-Aldrich, USA) and 40µM DCPIP (Sigma-Aldrich, USA), and assays were carried out in different concentration of Dicumarol (Calbiochem, USA). NQO1 activity is described as the dicumarol inhibitable decrease in absorbance at 600 nm with DCPIP as a substrate and is expressed in nanomoles of DCPIP reduced per minute per milligram of protein. Total protein in tissue preparations was determined by the Biorad Protein assay kit (Hercules, CA, USA) using bovine serum albumin as a standard.

### Single cell gel electrophoresis

For single cell gel electrophoresis or ‘COMET’ assay, individuals were asked to rinse their mouths thoroughly for 2min with tap water. The exfoliated buccal cells were collected from one or both cheeks from the controls, and in the area with lesion from the cases, depending on the region where the lesion was located, including cheek, soft and hard palate, dorsal, ventral and lateral surfaces of the oral cavity. Typically, sites with tough, leathery texture of mucosa, blanching of mucosa (persistent, white, marble-like appearance which may be localized, diffuse or reticular), quid-induced lesions (fine, white, wavy, parallel lines that do not overlap or criss-cross, are not elevated and radiate from a central erythematous area) were selected. Comet assay was performed and analyzed as described previously (Katarkar et al., 2014).

### Statistical analysis

The calculation of genotypic and allele frequencies for cases and controls and association between polymorphisms of NQO1 C609T with the risk of oral submucous fibrosis among total and stratified population was estimated by computing Odds ratio (OR) and calculating 95% Confidence Interval (CI) using a chi-square Table Analysis and Yates corrected P value was taken for significance. The expression of NQO1 protein was analysed densitometricaly using Image J software after proper calibrations. The difference in protein expression was analysed by Student “t” test, p value <0.05 was significant.

## Results

The study population was stratified according to Gender (Male and Female) and Median age (40 years) as presented in Table 1. We found that male (65.9%) constituted a greater percentage of the study sample and that most of the OSF cases were in the age group of > 40 years (65.9%).

**Table 1:** Demographic variables of the studied population.

|  | OSF cases (179) | Healthy Controls (152) |
| --- | --- | --- |
| **Age** |  |  |
| Range | 14-73 | 18-78 |
| Mean | 40 | 37 |
| **Gender** |  |  |
| Male | 118(65.9%) | 74 (48.6%) |
| Female | 61(34.1%) | 78 (52.4%) |
| **Above 40 yrs** | 118(65.9%) | 94 (61.8%) |
| **Below 40 yrs** | 61(34.1%) | 58 (39.1%) |

### Distribution of Genotypes of *NQO1 C609T* polymorphism among the cases and controls

Wild type C/C allele, heterozygous C/T allele and homozygous mutant T/T alleles were determined by PCR-RFLP method (Figure 1A). Sequencing of random samples were done to check the genotype and compared to those obtained by PCR-RFLP to rule out experimental errors. Distribution of genotypes among the study samples (cases and control) are presented in Table 2. Overall, the frequency of homozygous mutant type (18%) was significantly higher in OSF cases than controls (8.5%) [OR of 2.369 (1.167-4.804) P=0.026]. When stratified according to age, both the heterozygous carrier and homozygous mutant variant was found to be higher in cases above 40 years of age compared to controls OR 4.5 (1.77-11.61) P=0.003 and OR 6.4 (1.92-21.35) P=0.004 respectively.

**Table 2:** Genotypic distribution of NQO1 C 609 T polymorphism among cases and controls:

|  | **Cases** | **Normal** | **OR\*** | **95% CI\*\*** | **P Value** |
| --- | --- | --- | --- | --- | --- |
| **Total population** | n=179 | n=152 |  |  |  |
| Pro/Pro | 80 (44.7) | 77 (50.65) | ref |  |  |
| Pro/Ser | 67 (37.43) | 62 (40.78) | 1.056 | 0.664-1.679 | 0.913 |
| Ser/Ser | 32 (17.8) | 13 (8.5) | 2.369* | 1.167-4.804 | 0.026 |
| **Below 40 yrs** | n=128 | n=96 |  |  |  |
| Pro/Pro | 60 (46.87) | 33 (34.04) | ref |  |  |
| Pro/Ser | 48 (37.5) | 54 (56.3) | 0.493 | 0.27-0.87 | 0.024 |
| Ser/Ser | 20 (15.6) | 9 (9.5) | 1.185 | 0.49-2.85 | 0.893 |
| **Above 40 yrs** | n=51 | n=56 |  |  |  |
| Pro/Pro | 20 (39.2) | 43 (76.7) | ref |  |  |
| Pro/Ser | 19 (37.2) | 9 (16.07) | 4.5* | 1.77-11.61 | 0.003 |
| Ser/Ser | 12 (23.5) | 4 (7.1) | 6.4* | 1.92-21.35 | 0.004 |
| **Male** | n=118 | n=74 |  |  |  |
| Pro/Pro | 52 (44.06) | 30 (40.5) | ref |  |  |
| Pro/Ser | 46 (38.9) | 38 (51) | 0.698 | 0.376-1.297 | 0.329 |
| Ser/Ser | 20 (16.9) | 6 (8.1) | 1.923 | 0.711-5.59 | 0.3 |
| **Female** | n=61 | n=78 |  |  |  |
| Pro/Pro | 28 (45.9) | 32 (41.02) | ref |  |  |
| Pro/Ser | 21 (34.4) | 39 (50) | 0.615 | 0.297-1.276 | 0.265 |
| Ser/Ser | 12 (11.4) | 7 (8.9) | 1.95 | 0.95-5.15 | 0.322 |

\* Odds Ratio; \*\* Confidence interval

**Figure 1:**
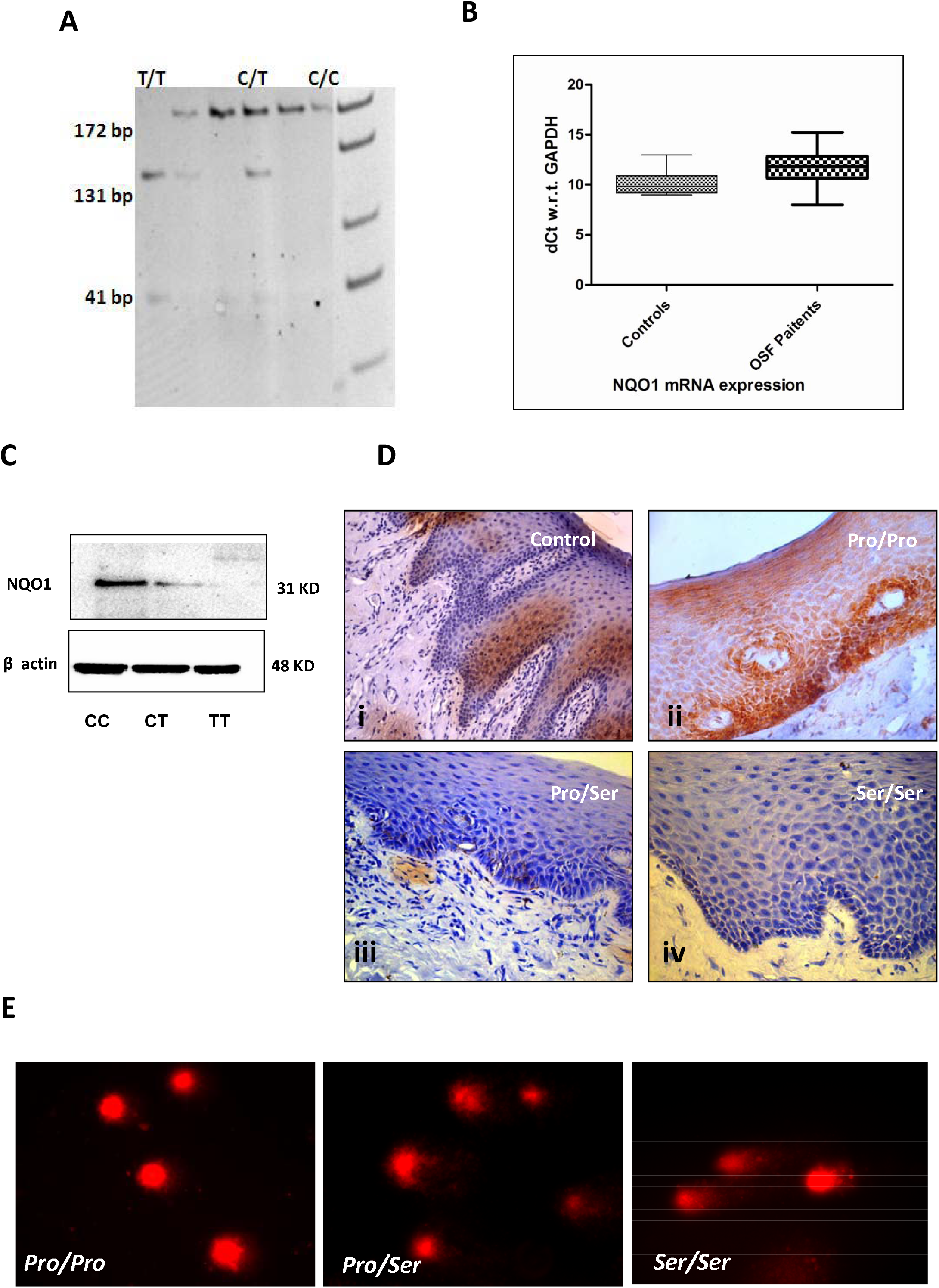
Effect of various genotypes on expression patterns of NQO1 and buccal tissue architecture. **A)** Genotype determination by restriction fragment length polymorphism following PCR amplification (PCR-RFLP). The size of digested products is shown in the figure. **B)** Increased mRNA expression *wrt* GAPDH of NQO1 in OSF cases compared to controls indicating transcriptional up regulation in affected tissues. **C)** Expression levels of NQO1 protein in different genotypic conditions in OSF cases. **D)** Immunohistochemical localization shows high basal level expression of NQO1 especially along the rete-ridges and submucosal cells lining epethilium in OSF cases with *Pro/Pro* allele (ii) compared to controls (i). Little or no detectable expression is observed in heterozygous (*Pro/Ser*) (iii) or homozygous (*Ser/Ser*) (iv) oral mucosal tissue. **E)** DNA damage as evident from ‘COMET’ assay performed on exfoliated buccal tissue from OSF cases with different genotypes. DNA disintegration worsens with heterozygous or homozygous mutant conditions.

### NQO1 expression and epethilial DNA disintegration in different genotypes

NQO1 mRNA expression level was higher in cases with either genotype compared to controls indicating transcriptional up regulation of NQO1 in tissues (Figure 1B). In general a significantly decreased mean expression for NQO1 protein was observed in buccal mucosal tissues of cases carrying heterozygous (42%) or homozygous (70%) genotypes (p=0.0055 & p=0.0001 respectively) indicating NQO1 degradation in homozygous / heterozygous mutant cases is a post-transcriptional event (Table 3) (Figure 1C).

**Table 3:** Effect of presence of NQO1 C609T polymorphism on protein expression (densitometric units) in OSF cases.

| Genotype |  |  | P value |
| --- | --- | --- | --- |
| *Pro/Pro Vs Pro/Ser* | *Pro/Pro* | *Pro/Ser* | 0.0055 |
|  | 1.4875±0.8046 | 0.8789±0.5969 |  |
| *Pro/Pro Vs Ser/Ser* | *Pro/Pro* | *Ser/Ser* | 0.0001 |
|  | 1.4875±0.8046 | 0.4297±0.2492 |  |

Histochemical localization of NQO1 protein in oral epithelial cells (Figure 1D) is concurrent with western-blots and shows high concentrations of NQO1 expression in epithelium of the tissue sections obtained from OSF cases with wild type *Pro/Pro* genotype compared to almost focal expression in normal healthy epithelium. Sections show increased NQO1 expression around the basal cell layer in wild type cases. NQO1 expression in heterozygous *Pro/Ser* OSF cases showed faint localizations whereas cases with homozygous mutant trait showed almost no NQO1 expression in the basal cells or elsewhere.

Comparative analysis of single cell gel electrophoresis of exfoliated buccal cell collected from the site of the lesion, reveals higher degree of disintegrated DNA in heterozygous and homozygous mutant OSF cases (Figure 1E) compared to wild types.

### Lower NQO1 activity in OSF and OSF associated with malignancy tissues compared to their adjacent tissue

Figure 2A presents kinetics of NQO1-mediated reduction of DCPIP as a measure of NQO1 activity. The Vmax and Km of NQO1 at different concentration of dicoumarol inhibitor in Normal, OSF and OSF associated with malignancy cases are presented in supplementary table 1. With increase in concentration of Dicoumarol (0, 1.25, 2.5, 5 and 10 µM) there was a uniform decrease in Vmax of NQO1 but no significant change in Km value was detected either in presence or absence of the activator (BSA). This suggests there is a non-competitive inhibition of the NQO1 activity in the presence of inhibitor. When compared to the OSF tissue alone there was about 2-fold decrease in activity (p=0.028) with or without BSA (supplementary figure 1). This was about 10-fold decreased in OSF associated with malignant tissues (p< 0.0001) thus explaining insignificant NQO1 activity.

**Figure 2:**
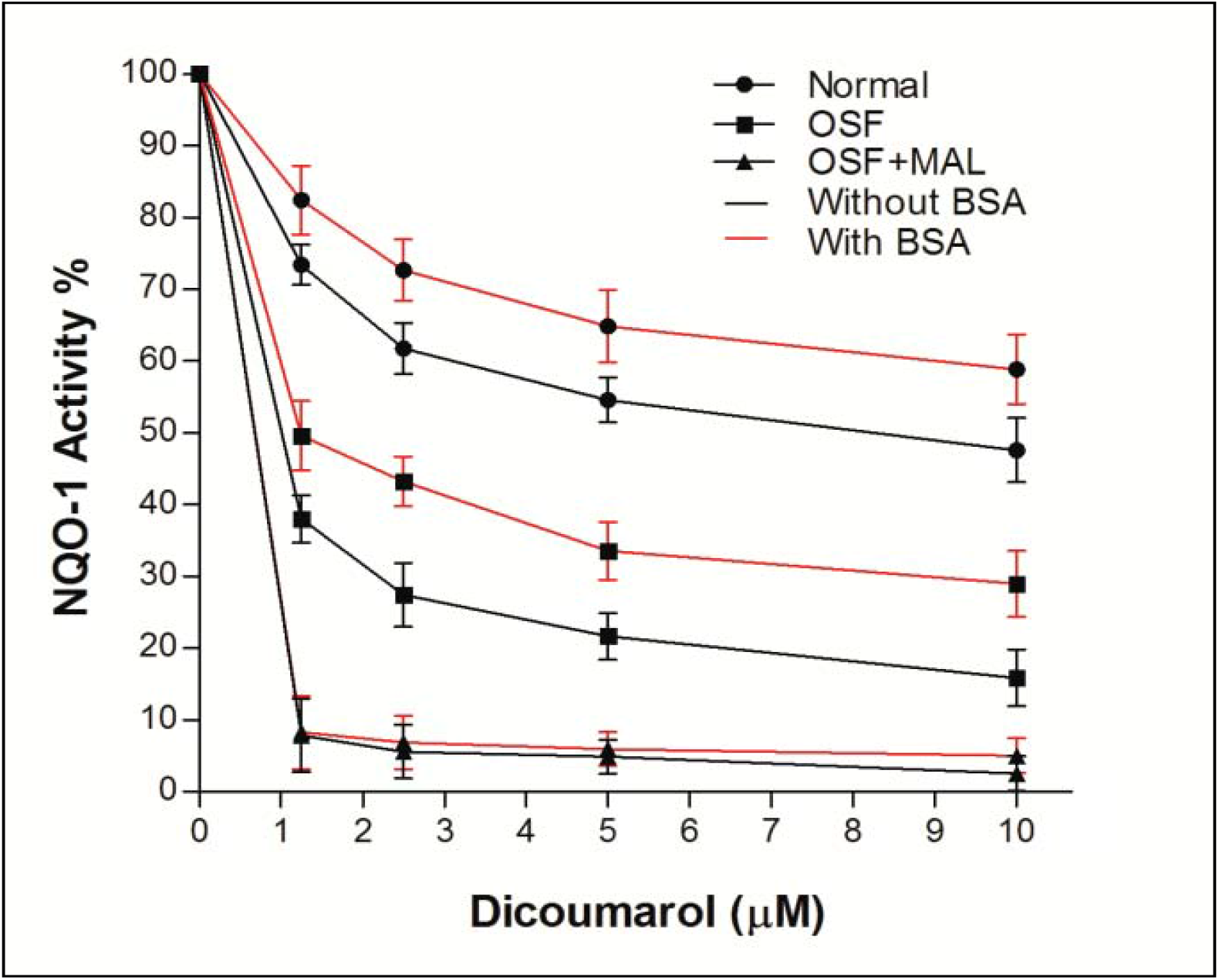
NQO1 activity as measured by reduction of DCPIP in presence of dicoumarol with or without activator. The OSF with malignant lesion cases (OSF+MAL) carrying Ser/Ser allele presents reduced or no activity of NQO1 compared to normal or OSF cases with no malignancy.

## Discussion

Oral submucous fibrosis is a potentially malignant condition affecting any part of the oral cavity and pharynx, which could subsequently develop into oral cancer (Arora et al., 2014). The carcinogenic turnover of OSF may serve as a very important model to study various molecular changes that is taking place during oral carcinogenesis. In the present study we have tried to understand the pathogenesis and carcinogenesis of OSF in the light of NQO1 C609T polymorphism. In OSF mucosal tissues from cases carrying *Ser/Ser* genotype, either traces or no NQO1 protein was detected, this might be due to ubiquitin mediated proteasomal degradation as discussed earlier (Siegel et al., 2001). In the present study, we observed a significantly higher number of subjects carrying the NQO1 homozygous mutant allele genotype in cases suffering from oral submucous fibrosis as compared to the control group. Mostly male cases with OSF reported to our outpatient department (OPD) compared to the female cases with a median of 40years of age. Cases above 40yrs showed a greater association with the risk of the disease. This result is in line with previous investigations that showed an association between the NQO1 homozygous mutant genotype and other tumour types such as urothelial cancer, renal cancer, leukemias, lung cancer and cutaneous basal cell carcinoma (Liu and Zhang; Schulz et al., 1997; Clairmont et al., 1999; Guha et al., 2008; Wang et al., 2008). In addition, we also observed that NQO1 protein expression at the site of the lesion was reduced (42% in *Pro/Ser* and 70% in *Ser/Ser*). The mechanism underlying the correlation of NQO1 C609T polymorphism with the increased risk for developing various tumours likely resides in the different enzyme activities encoded by the NQO1 alleles.

The current study suggests that the minor T allele acts in a recessive mode in the development of OSF, demonstrated by the fact that the heterozygous genotype frequency among OSF cases remains similar to that among the healthy controls. This indicates that in individuals with the NQO1 heterozygous genotype, the enzyme activity might be enough for protecting cells from damage by exogenous carcinogens important for the development of OSF. One of the major limitations in our study was availability of biopsy tissues especially from OSF associated with malignancy cases for any further functional studies, that could have been more suggestive of the mechanism of carcinogenesis.

The determination of the NQO1 C609T genotype may be valuable as a stratification marker in future intervention trials for OSF and oral squamous cell carcinoma (OSCC). This finding may be particularly important in our country as most of the common people are habituated to areca chewing in different modes and are susceptible to development of OSF and eventually OSCC. Several cancer chemotherapeutic drugs have been developed targeting NQO1 (Oh and Park, 2015). Since NQO1 C609T polymorphism is found to be positively associated with many solid as well as blood malignancies, therefore, a practical approach for cost-effective tumour screening may be designed taking other such gene polymorphisms, especially the metabolic genes (Chaudhuri et al., 2013) into account before starting a treatment regimen.

## Acknowledgement

The study was supported by Council of Scientific and Industrial Research, Govt. of India. SM is grateful to CSIR for Senior research Fellowship. AK is grateful to ICMR for Senior research Fellowship

## Figure Legends

**Supplementary Figure 1:** Graphical presentation of NQO1 activity in presence of dicoumarol in normal, OSF and OSF with malignancy (OSF+MAL) tissue extracts in absence (gray bars) or presence (black bars) of BSA as activator.

