## Supplementary material for "Association of NQO1 C609T (Pro187Ser) with risk of Oral Submucous Fibrosis in Eastern Indian population": Supplemetary table 1

|  | Normal | | | |
| --- | --- | --- | --- | --- |
| Dicoumarol (µM) | With BSA | | Without BSA | |
|  | Vmax | Km | Vmax | Km |
| 0 | 104.9 ± 9.17 | 4.83 ± 1.29 | 84.72 ± 7.14 | 4.63 ± 1.21 |
| 1.25 | 103 ± 8.28 | 5.57 ± 2.28 | 76.64 ± 3.70 | 5.50 ± 2.76 |
| 2.5 | 87.16 ±7.11 | 5.72 ± 1.32 | 75.18 ± 5.10 | 8.98 ± 1.42 |
| 5 | 61.61 ± 6.94 | 5.74 ± 1.83 | 21.98 ± 3.07 | 5.47 ± 2.21 |
| 10 | 37.55 ± 3.11 | 3.56 ± 2.03 | 15.73 ± 2.53 | 4.58 ± 2.31 |
|  | OSF | | | |
| Dicoumarol (µM) | With BSA | | Without BSA | |
|  | Vmax | Km | Vmax | Km |
| 0 | 37.48 ± 3.27 | 5.31 ± 2.24 | 20.55 ± 1.74 | 3.63 ± 2.12 |
| 1.25 | 35.52 ± 2.85 | 6.67 ± 1.43 | 16.86 ± 2.81 | 5.32 ± 2.89 |
| 2.5 | 26.95 ± 2.2 | 5.49 ± 2.53 | 17.76 ± 1.26 | 5.83 ± 1.93 |
| 5 | 17.14 ± 1.93 | 4.93 ± 1.57 | 4.78 ± 2.66 | 5.11 ± 2.17 |
| 10 | 9.57 ± 0.79 | 3.41 ± 1.63 | 3.19 ± 3.14 | 4.14 ± 1.37 |
|  | OSF+MAL | | | |
| Dicoumarol (µM) | With BSA | | Without BSA | |
|  | Vmax | Km | Vmax | Km |
| 0 | 7.47 ± 1.61 | 2.61 ± 2.25 | 7.05 ± 1.58 | 2.46 ± 2.26 |
| 1.25 | 6.78 ± 1.44 | 2.77 ± 2.29 | 6.21 ± 1.27 | 2.48 ± 2.08 |
| 2.5 | 5.57 ± 1.61 | 2.83 ± 3.13 | 5.01 ± 1.59 | 2.77 ± 3.43 |
| 5 | 3.41 ± 0.56 | 2.01 ± 1.48 | 3.07 ± 0.57 | 2.27 ± 1.8 |
| 10 | 2.31 ± 0.34 | 2.24 ± 1.41 | 1.98 ± 0.27 | 2.77 ± 1.47 |

**Supplementary Table 1: Vmax and Km of NQO-1 at different concentration of dicoumarol inhibitor in Normal, OSf and OSF+Mal patient .**
