## Supplementary figures and images for "Association of NQO1 C609T (Pro187Ser) with risk of Oral Submucous Fibrosis in Eastern Indian population"

### Supplementary table 2

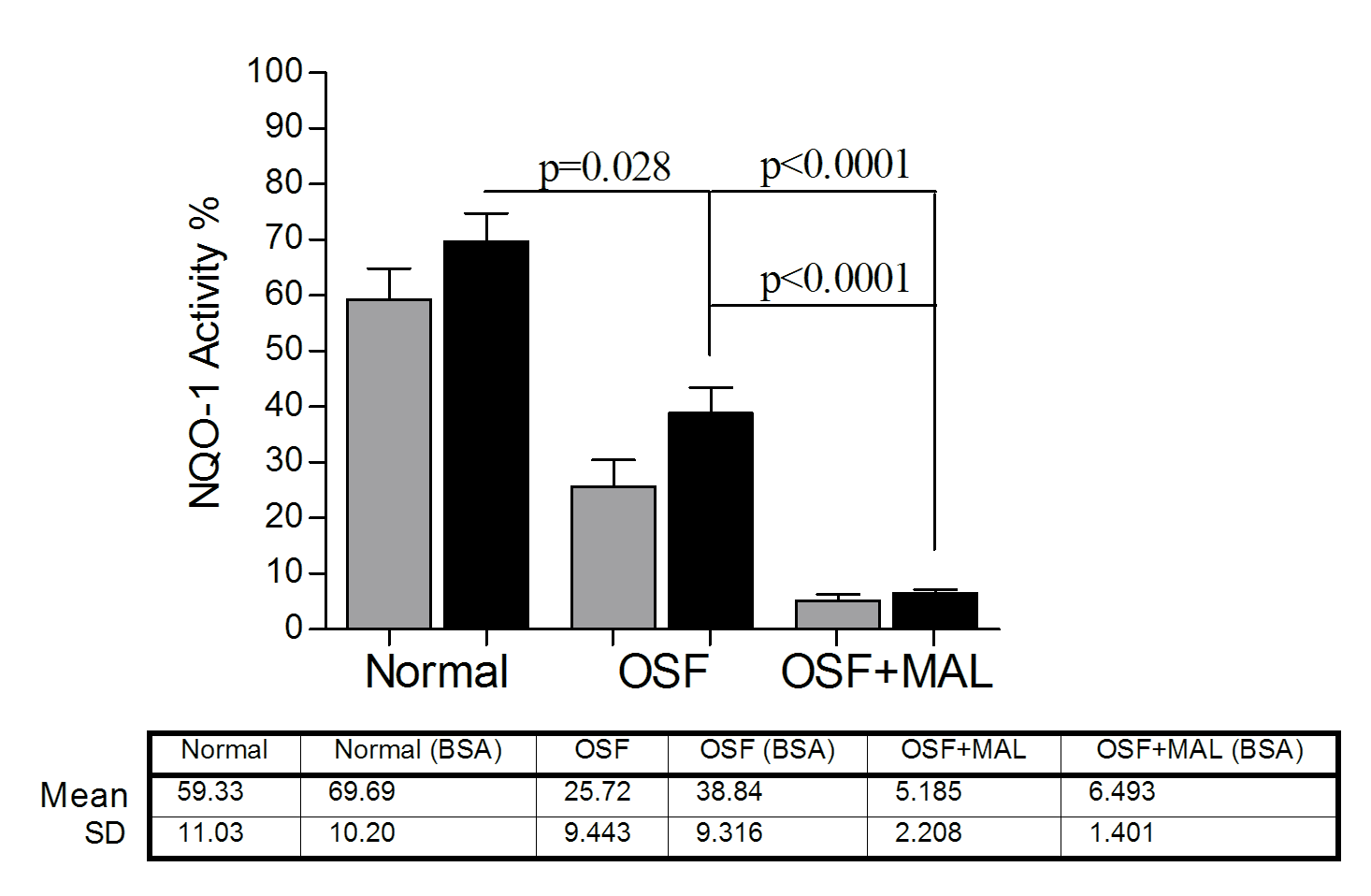
